## Supplementary information for "Topography-induced large-scale anti-parallel collective migration in vascular endothelium"

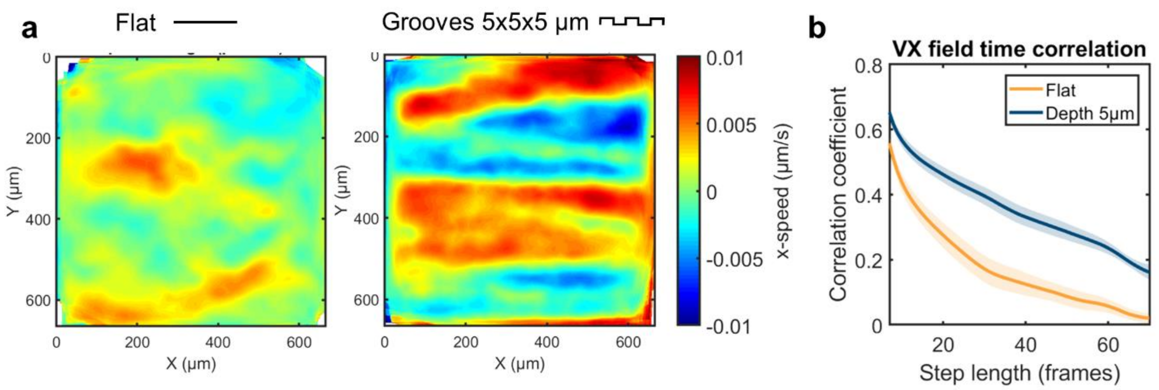


**Supplementary Figure 1: Establishment of the stream pattern on microgrooved substrates. a**, Time-averaged x-direction velocity fields on flat surfaces and on 5 x 5 x 5 µm (w x s x d) grooves. **b**, Quantification of the time correlation length: the Pearson correlation coefficient is calculated for pairs of frames increasingly distant in time. Error bars represent SEM, n=3 independent experiments.

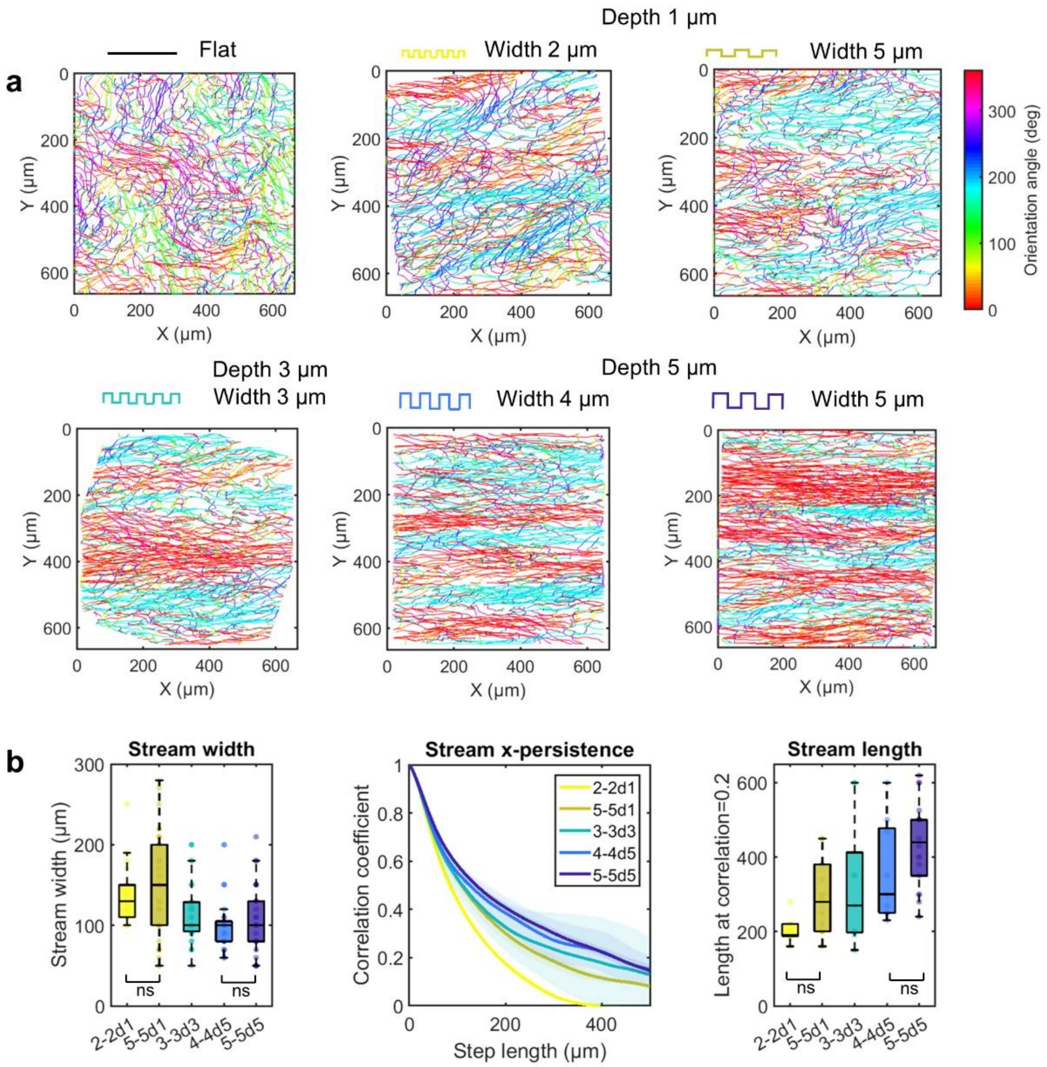


**Supplementary Figure 2: Influence of groove width on the spatial features of the streams. a**, Accumulated cell trajectories after 24 h of migration for flat control substrates and for microgrooved substrates of different dimensions: w=s=2 µm d=1 µm (2-2d1), w=s=5 µm d=1 µm (5-5d1), w=s=3 µm d=3 µm (3-3d3), w=s=4 µm d=5 µm (4-4d5), w=s=5 µm d=5 µm (5-5d5). Trajectories are color-coded for the orientation angle of each displacement vector. **b**, Quantification of the spatial features of the streams with the same method as explained in the main text. One-way ANOVA, Dunn's/Fisher's post-test, n=3 independent experiments.

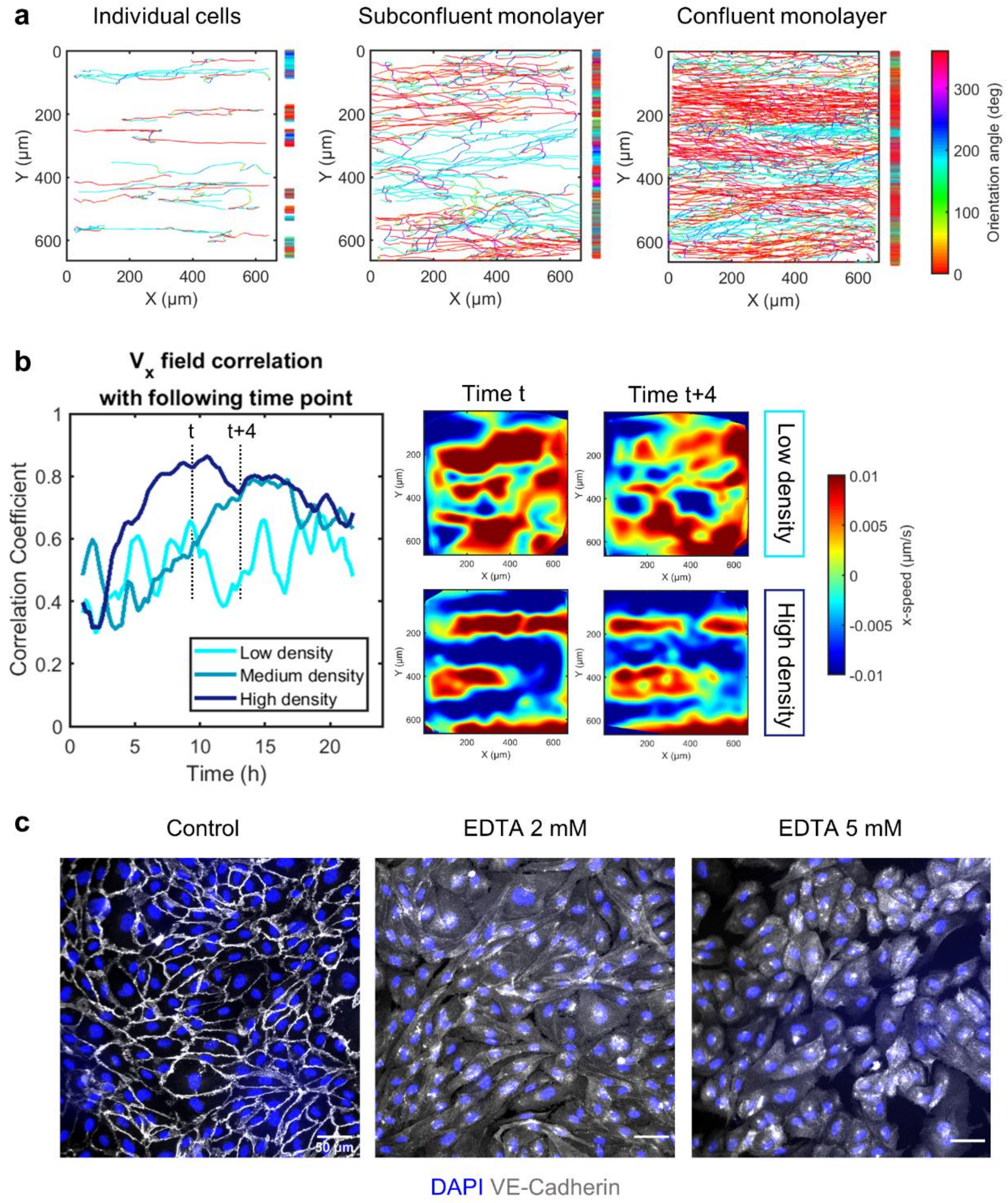


**Supplementary Figure 3: Emergence of cell streams with increasing cell density. a**, Accumulated cell trajectories after 24 h of migration for individual cells, low-density monolayer, and high-density monolayer on grooves of width, spacing, and depth of 5 µm. Trajectories are color-coded for the orientation angle of each displacement vector, and vertical bars represent the projected trajectories on the y-axis. **b**, Time-averaged x-direction velocity field for monolayers of different densities and immunostaining against VE-cadherin showing the morphology of cell-cell junctions for the approximate same cell density. Scale bar, 30 µm. The fields of view (FOVs) correspond to movie3. The graph shows for these 3 FOVs the time evolution of the correlation between the $V_x$ field at time t and at t+1 and illustrates the dynamic evolution of the collective pattern of motion. **c**, Immunostaining against VE-cadherin and DAPI on HUVEC monolayers after 48 h of culture on flat surfaces for control, 2 mM EDTA, or 5 mM EDTA treatment (24 h).

**Supplementary note: Theoretical modeling**

We explain the emergence of endothelial cell streams by an active fluid model inspired by that proposed by Duclos *et al.*[3]. Duclos *et al.* applied the model of Voituriez *et al.*[18] based on active gel theory [19] to describe spontaneous flows in confined cell sheets. They explained the emergence of global directional motion of otherwise non-migrating cells in a confined stripe with cells advancing in opposite directions at the two ends of the stripe. To account for the specificities of our experiment, the model differs in two main aspects from that by Duclos *et al.* First, an additional term in the free energy equation accounts for the tendency of cells to orient along the ridges. Second, an active term is included to account for single cell locomotion, in addition to the cell contractility term already accounted for by Duclos *et al.*

*Active fluid equations.* We consider a 2D coordinate system $(x,y)$ with the $x$-direction aligned with the ridges and the $y$-direction perpendicular to them. Endothelial cells are treated as active nematic particles whose orientation is characterized by the unit director field $\boldsymbol{p} = (\cos\theta, \sin\theta)$, with $\theta$ being the angle between the cell orientation and the ridges. Because endothelial cells are able to change the direction of motion without rotating the cell body, we model them as nematic particles. Actomyosin cell activity induces a contractile (or extensile) force dipole acting on each cell. The force per unit length of the cell is given by $\zeta\Delta\mu$, where $\Delta\mu$ is the free energy produced by nutrient consumption and $\zeta$ is the reactive coefficient as defined in active gel theory[19], whose sign indicates whether active forces are contractile ($\zeta<0$) or extensile ($\zeta>0$). Note that the term $\zeta\Delta\mu$ is a force dipole. Therefore, the associated net force acting on a single cell is zero, and this term does not lead to single cell locomotion. To account for single cell locomotion, we account for an additional active term consisting of a net active force that acts along the cell orientation. This additional active force corresponds to the reaction force from the substrate due to cell contractility coupled with substrate adhesion at the cell front (or equivalently cell extension coupled with adhesion at the rear).

*Free energy*. The free energy of the director field in two dimensions is given by

$$\mathcal{F} = \int \left(\frac{K_1}{2}(\nabla \cdot \boldsymbol{p})^2 + \frac{K_3}{2}(\nabla \times \boldsymbol{p})^2 + \frac{\alpha}{2}p_y^2\right). \tag{1}$$

The first two terms are the classic formulation of the free energy in active gels, where $K_1$ and $K_3$ are the splay and bend Frank constants, respectively. As in Duclos *et al.*, we assume $K_1 = K_3 = K$. The third term is a new term that we introduce to penalize cell orientation across the ridges. The coefficient $\alpha$ is a measure of the tendency of the cells to align with the ridges that is expected to depend on the ridge height. By using the Einstein relation between diffusion and active thermal agitation, this coefficient can be estimated as $\alpha \approx \xi_\mathrm{x} D_\mathrm{t}/\sigma_\theta^2$, where $\xi_\mathrm{x}$ is the substrate friction coefficient, $D_\mathrm{t}$ the translational diffusion, and $\sigma_\theta$ the standard deviation of cell orientation angles. We have measured for our cells $D_\mathrm{t} \approx 10$ µm$^2$/min, which leads us to estimate $\alpha$ to be in the range $10^{-3}$–$10^{-2}$ J/m$^2$, depending on the value of $\sigma_\theta$ which is determined by the groove geometry.

*Constitutive relations*. The linearized constitutive relations for an active nematic fluid in two dimensions relate the stress $\boldsymbol{\sigma}$ to the cell orientation $\boldsymbol{p}$ as well as the variations in cell orientation to the shear stress. They are:

$$\sigma_{\alpha\beta} = 2\eta u_{\alpha\beta} - \zeta\Delta\mu\, p_\alpha p_\beta + \frac{\nu}{2}\left(p_\alpha h_\beta + p_\beta h_\alpha\right) + \frac{1}{2}\left(p_\alpha h_\beta - p_\beta h_\alpha\right) - P\delta_{\alpha\beta}, \tag{2}$$

$$\partial_t p_\alpha + v_\beta \partial_\beta p_\alpha + \omega_{\alpha\beta} p_\beta = \frac{h_\alpha}{\gamma} - \nu u_{\alpha\beta} p_\beta. \tag{3}$$

Here, $\alpha$ and $\beta$ are indices that stand for the directions $x$ or $y$, with a repeated index implying summation over all values of the indices, as per Einstein's convention. $\sigma_{\alpha\beta}$ is the stress tensor, $u_{\alpha\beta} = (\partial_\alpha v_\beta + \partial_\beta v_\alpha)/2$ is the shear rate, with $v_\alpha$ the velocity component, and $\omega_{\alpha\beta} = (\partial_\alpha v_\beta - \partial_\beta v_\alpha)/2$ is the vorticity tensor. $P$ is a Lagrange multiplier, related to the pressure, that enforces incompressibility, i.e., $\partial_\beta v_\beta = 0$. $h_\alpha = -\partial\mathcal{F}/\partial p_\alpha$ is the so-called orientational field. $\eta$ and $\gamma$ are the shear and rotational viscosities, respectively, arising from both cell mechanical properties and cell-cell interactions. υ is the flow alignment parameter, which controls how mechanical stress determines cell orientation.

*Force balance*. The force balance equation, assuming negligible inertia, reads

$$\partial_\beta \sigma_{\alpha\beta} = \xi_\alpha \left( v_\alpha - v_{0,\alpha} \frac{v_\alpha}{|v_\alpha|} \right), \tag{4}$$

where $\xi_\alpha$ is the friction coefficient along the $\alpha$-direction. Duclos *et al.* concluded that substrate friction did not significantly change the physics of the phenomenon they considered. Here we will show that substrate friction is needed to explain the emergence of periodic cell streams. Because in our setup individual cells migrate, Eq. (4) includes the term $v_{0,\alpha}$, which corresponds to the active migration velocity of a single cell. For simplicity, we will assume $v_{0,x} = v_0$ and $v_{0,y} = 0$.

*Simplification of the equations*. We seek a quasi-1D solution to the problem defined by Eqs. (1-4) by assuming the solution to be uniform along *x*. Moreover, we assume that the velocity along the ridges $v_x$ dominates the transverse velocity $v_y$, which we neglect. We express the orientational field by its components parallel and perpendicular to the director, $h_\perp$ and $h_\parallel$. We obtain:

$$h_\perp = \gamma \left[ \frac{(\nu \cos 2\theta + 1)}{2} \frac{\partial v_x}{\partial y} + \frac{\partial \theta}{\partial t} \right], \tag{5}$$

$$h_\parallel = \frac{\gamma\, \nu \sin 2\theta}{2} \frac{\partial v_x}{\partial y}. \tag{6}$$

Next, we linearize the force balance equation in the *x*-direction, Eq. (4), by assuming $\theta \ll 1$, to obtain:

$$\left(\eta + \frac{\gamma(1+\nu)^2}{4}\right)\frac{\partial^2 v_x}{\partial y^2} - \zeta\Delta\mu\frac{\partial\theta}{\partial y} + \frac{\gamma(1+\nu)}{2}\frac{\partial^2\theta}{\partial t\partial y} = \xi_x\left(v_x - v_0\frac{v_x}{|v_x|}\right). \qquad (7)$$

By using Eq. (5) and the relationship between the orientational field and the free energy, $h_\alpha = -\partial\mathcal{F}/\partial p_\alpha$, we obtain:

$$K\frac{\partial^2\theta}{\partial y^2} - \alpha\theta = \frac{\gamma(1+\nu)}{2}\frac{\partial v_x}{\partial y} + \gamma\frac{\partial\theta}{\partial t}. \qquad (8)$$

Eqs. (7) and (8) constitute a system of two ODEs with two unknowns: $v_x(y,t)$ and $\theta(y,t)$.

*Linear stability analysis.* We perform a linear stability analysis of the system formed by Eqs. (7) and (8) around an equilibrium state where cells would all be moving along the +*x* ridge direction, $\theta_{\mathrm{eq}}(y)=0$, at the single-cell velocity, $v_{x,\mathrm{eq}}(y)=v_0$. We are specifically interested in finding the wavelength of the fastest growing mode, which will indicate the expected spatial periodicity of the flow profile. Because the perturbation is small, $v_x>0$. Thus, in Eq. (7) we can make $v_x/|v_x|=1$. We obtain the following dispersion relationship between the rate of growth σ and the wavenumber *k*:

$$\sigma = -\frac{(\alpha + Kk^2)}{\gamma}\left(1 + \frac{\gamma(1+\nu)^2k^2}{4(\xi_x + \eta k^2)}\right) - \frac{\zeta\Delta\mu(1+\nu)k^2}{2(\xi_x + \eta k^2)}. \qquad (9)$$

In the limit $\alpha=0$, this equation is identical to that obtained by Duclos *et al.* in the presence of substrate friction. Unstable modes can arise if $\zeta\Delta\mu(1+\nu) < 0$, for example in the case of contractile cells ($\zeta\Delta\mu < 0$). We can show that a threshold wavelength exists below which the system is unstable. The most unstable wavenumber is:

$$k_* = -\frac{\xi_x}{\eta} + \sqrt{\left(\frac{\xi_x}{\eta}\right)^2 + \frac{1}{l^4}}, \qquad (10)$$

where

$$l^4 = \frac{K\eta\left[\frac{4\eta}{\gamma} + (1+\nu)^2\right]}{\xi_x\left[2(-\zeta\Delta\mu)(1+\nu) - \alpha(1+\nu)^2 - \frac{4K\xi_x}{\gamma}\right]}. \quad (11)$$

For $k_*$ to exist, the term in square brackets in the denominator of the right-hand side of Eq. (11) must be positive, which requires $2(-\zeta\Delta\mu)(1+\nu) > \alpha(1+\nu)^2$, since according to our parameter estimates, the third term in the denominator containing $K$ is small compared to the other two. Thus, we assume the bracketed term in the denominator to be of the order of the active contractile term. With our parameter estimates (Table 1), we obtain $l^4 \approx 2.5 \cdot 10^3$ μm$^4$ and $k^{*2} \approx 2 \cdot 10^{-2}$ μm$^{-2}$, which corresponds to a stream width $W^* = 2\pi/k^* \approx 50$ μm.

Table 1: Estimates of model parameters.

| Symbol | Meaning | Value | Units |
|---|---|---|---|
| $\lvert -\zeta\Delta\mu \rvert$ | Cell contractility | $10^{-2}$ | Pa·m |
| $\eta$ | 2D shear viscosity | 10 | Pa·s·m |
| $\gamma$ | 2D rotational viscosity | 10 | Pa·s·m |
| $\lvert 1+\nu \rvert$ | Flow alignment parameter | 1 | |
| $\xi_x$ | Substrate friction coefficient | $10^{10}$ | Pa·s·m$^{-1}$ |
| $K$ | Frank constant | $10^{-14}$ | Pa·m$^3$ |
| $\alpha$ | Ridge alignment constant | $10^{-3}$–$10^{-2}$ | Pa·m |
| $v_0$ | Single cell migration speed | $10^{-8}$ | m·s$^{-1}$ |

*Equilibrium periodic velocity profile.* The linear stability analysis above suggests the emergence of streams. We now consider the equilibrium hydrodynamics. We note that active cell propulsion will require that cells flow with velocities of the order of $v_x \approx \pm v_0$, whereas continuity in the number of cells requires zero average velocity over the width. Based on these conditions, we investigate whether a velocity profile with alternating streams satisfies the governing equations and, if so, what is the required width of the streams.

We thus assume a velocity profile that is periodic along $y$, with period defined as $2W$, where $W$ is the constant stream width to be determined. Mathematically, this periodicity is described by $v_x(y+2W)= v_x(y)$ and $\theta(y+2W)= \theta(y)$. Symmetry between the $+x$ and $-x$ directions suggests that streams of opposite direction have an equal width of $W$, with the corresponding velocity profiles being equal and opposite. We chose the origin $y$=0 such that $v_x$>0 for $-W/2<y< W/2$, and we solve the governing equations (7) and (8) for a steady-state profile. Based on the results of the linear stability analysis and on parameter estimates, we expect the term containing the second derivative in Eq. (8) to be small compared to the other two, which yields the relationship

$$\theta \approx -\frac{\gamma_e}{\alpha}\frac{\partial v_x}{\partial y}, \tag{12}$$

where $\gamma_e = \gamma(1+\nu)/2$. Substituting this relationship into Eq. (7), we obtain an ordinary differential equation for the velocity profile:

$$\beta\frac{\partial^2 v_x}{\partial y^2} + \xi_x v_x = \xi_x v_0\,, \tag{13}$$

where $\beta = (\gamma_e/\alpha)(-\zeta\Delta\mu) - \eta_e$ and $\eta_e = \eta + \gamma(1+\upsilon)^2/4$. According to our parameter estimates, the two terms in the expression of $\beta$ are of comparable magnitude. Two cases are therefore possible: $\beta$<0 and $\beta$>0. The case $\beta$<0 leads to a periodic velocity profile described by hyperbolic functions. However, free energy minimization for this solution corresponds to

$W$=0 or $W\to\infty$, inconsistent with experimental observations. We thus postulate that the current experiments belong to the strong contractility regime for which $\beta>0$. In this regime, we solve Eq. (12) subject to the boundary conditions $v_x=0$ at $y=\pm W/2$. Among all possible solutions, we choose the solution that minimizes the free energy, Eq. (1). Because $K/W^2 \ll \alpha$, minimizing the free energy corresponds to minimizing the average value of $\theta^2$. We thus obtain the equilibrium velocity profile

$$v_x = v_0 \left[1 + \cos\left(\pi \frac{2y}{W}\right)\right] \tag{14}$$

for $-W/2 < y < W/2$, and equal and opposite in the interval $W/2 < y < 3W/2$. The value of $W$ that minimizes the free energy of the system is given by

$$W = \frac{2\pi}{\sqrt{\xi_x}} \left[\frac{(1+\nu)}{2\alpha}\gamma(-\zeta\Delta\mu) - \eta - \frac{\gamma(1+\nu)^2}{4}\right]^{1/2}. \tag{15}$$

With the parameter estimates reported in Table 1, and for the case $\beta>0$, we can estimate the term in the square brackets to be of the order of the contractility term. This leads to an estimate of the stream width $W \approx 130$ µm.

According to the model, the maximum deviation of the cell orientation with respect to the ridges is $\theta_{\max} = \frac{v_0\gamma(1+\nu)\pi}{\alpha W} \approx 0.22$ rad, consistent with observations and supporting the hypothesis of small $\theta$ used in deriving this solution.

*Length of the streams*. We postulate that varying the ridge height modifies the length of the streams by modifying the alignment constant $\alpha$. When cells are placed on the ridges, their orientation will be stochastic and expected to follow a Boltzmann distribution, i.e. a normal distribution of zero mean and standard deviation given by

$$\sigma_\theta = \sqrt{\frac{k_\mathrm{B} T_\mathrm{eff}}{\alpha a^2}}, \tag{16}$$

where $a$ is the cell size. At a fixed time, the offset along the $y$ axis between two consecutive cells along a stream can be described as a one-dimensional random walk. The average step of this random walk is given by $\delta_y = \langle|\theta|\rangle a$. After $N$ steps, the length of the stream will be $L \approx Na$, whereas its width will be $\langle W^2\rangle^{1/2} = \delta_y\sqrt{N} \approx \sigma_\theta\sqrt{La}$. We can define the persistence length of the stream as the value of $L$ for which the offset $W$ becomes comparable to the cell length, $\langle W^2\rangle^{1/2} \approx a$, which yields

$$L \approx \frac{a}{\sigma_\theta^2} = \frac{\alpha a^3}{k_\mathrm{B} T_\mathrm{eff}}. \tag{17}$$
